# Generating and testing reagents for CRISPR/Cas9 based homologous recombination and gene drive in *Tribolium*

**DOI:** 10.1101/2023.11.07.566100

**Authors:** Hannah C. Markley, Kennedy J. Helms, Megan Maar, Gabriel E. Zentner, Michael J. Wade, Andrew C. Zelhof

## Abstract

CRISPR/Cas9 gene drive systems are possible in a few insects and ever expanding. Nonetheless, success in one species and techniques developed for it are not necessarily applicable to other species. As such, the development and expansion of gene drive systems is dependent upon direct experimentation. A critical aspect and potentially limiting factor of gene drive is the ability to induce Cas9-dependent homologous recombination. Here we report our attempts to induce Cas9-dependent homologous recombination and subsequent gene drive in *Tribolium castaneum*. Utilizing constructs containing one or two target gRNAs in combination with Cas9 under two different promoters and corresponding homology arms, we found a high incidence of CRISPR/Cas9 induced mutations but a complete lack of evidence of homologous recombination and genetic drive. Even though the generated constructs provide new resources for CRISPR/Cas9 modification of the *Tribolium* genome, our results suggest that *Tribolium* genome may be refractory towards Cas9-induced homologous recombination and additional modifications will be necessary to increase the potential for homologous recombination.

## Introduction

CRISPR/Cas9 gene drive systems have increased the possibility of devising synthetic methodologies, versus utilizing naturally occurring selfish elements, for controlling insect pests and limiting the spread of disease vectors [1-3]. CRISPR based gene drives have now been demonstrated in *Drosophila* and mosquitos, both *Anopheles* and *Aedes* [4-8], but success and ability to drive through a population is dependent on numerous factors (e.g. diploid versus diploid/haploid genomes [9], nature (e.g., sterility, viability, maternal effect) and timing of drive-gene expression [10], mating system [11], and frequency of drive-resistant alleles [11, 12]. Moreover, CRISPR/Cas9 drive systems are dependent on the choice of DNA double-strand break repair via homologous recombination versus non-homologous end joining (NHEJ). Homologous directed repair (HDR) is necessary to permit the conversion of a targeted locus and thus subsequently change inheritance frequencies. Whereas, if NHEJ occurs, it tends to be error prone, creating by insertion or deletion new alleles which are refractory to a gene drive system. Moreover, HDR is restricted to the S and G2 phases of the cell cycle while NEJH is active throughout the cell cycle [13]. Therefore, CRISPR/Cas9 drive based mechanisms are not only dependent on the molecular mechanisms of repair but also unique aspects of the species of interest.

*Tribolium castaneum*, like *Drosophila*, is an established model organism for developmental, evolutionary, and applied (e.g., insecticide) biology but is also a known agriculture pest throughout the world [14-20]. Hence the ability to demonstrate CRISPR/Cas9 gene drive and to a lesser extent a proven methodology for efficiency homologous directed repair would be beneficial to the research community. The genetic tools available to *Tribolium* include transgenesis, RNAi, and CRISPR/Cas9 gene editing [18]. Cas9 editing has been achieved by injection of ribonucleoprotein (RNP) complexes [21-23], plasmid encoded reagents [24-26], and injection of plasmid encoded gRNAs into transgenic *Tribolium* expressing Cas9 [27]. CRISPR/Cas9 edited genes include but not limited to *vermilion* [21, 27], *E-cadherin* [24], *cardinal* [22, 23], and *foxQ2* [28] as well as the ability to target inserted exogenous sequences like GFP [24, 25]. Whereas the capability to utilize CRISPR/Cas9 edit the genome is established in *Tribolium*, the extent and possibility of homologous directed repair is limited to only two reports and with very low frequency [26, 29]. Here we attempted to demonstrate the ability of CRISPR/Cas9 gene drive in *Tribolium*. Our results indicate that even though genome editing was achieved, our injections did not result in any recovered HDR modifications, only NHEJ repair. Subsequently we could not test for gene drive and thus possibly subsequent methodologies will be needed to increase a bias towards homologous directed repair in *Tribolium*.

## Materials and Methods

Tribolium husbandry and strains: All animals were raised at 28°C on a standard flour yeast mix. The following strains were utilized: *vermilion*^*white*^ (*v*^*w*^), [30], GA-1, and Henderson Black (HB) [31].

### Vectors and gRNA sequences

#### Tc-v 1gRNA backbone drive homology construct

The following components were synthesized and cloned into pUC57 (Synbio Technologies). *vermilion* left homology arm, U6a promoter driving the expression of Tcv95 gRNA [21], AscI cloning site, and *vermilion* right homology arm.

#### Tc-v 2gRNA backbone drive homology construct

The following components were synthesized and cloned into pUC57 (Synbio Technologies). *vermilion* left homology arm, U6a promoter driving the expression of TcV95 gRNA (5’-AAATTAAGTGAAGCCCAAGAAGG-3 ‘) [21], U6b promoter driving expression of TcV412 gRNA (5’-GGATCAAAACAACACGATTGAGG-3’), AscI cloning site, and *vermilion* right homology arm.

*hsp68-nls-Cas9-nls-hsp3’UTR* cassette was excised from p(bhsp68-Cas9) [24] Addgene (#65959) using flanking Asc1 sites and ligated into the AscI of both 1 or 2 gRNA drive homology constructs.

*nanos-nls-Cas-9-nls-T2A-EGFP-nanos UTR* cassette was flanked by AscI sites synthesized and cloned into pUC57 (Synbio Technologies). The *nanos* promoter consists of 347 bp of the first coding Methionine of *nanos* and *nanos* 3’UTR sequence is represented by 406 bp of DNA immediately downstream of the *nanos* termination codon. The Cas9 cassette was excised using flanking Asc1 sites and ligated into the AscI of both drive homology constructs.

Tribolium CRISPR injections: Injections were performed at 25°C and embryos were then returned to 28°C for development and hatching. Each construct was resuspended in 1x injection buffer (0.5 mM KCl; 0.01 mM NaPO4 buffer pH 7.5) at a concentration of 1 μg/μL and injected into GA-1 or Henderson Black (HB) embryos. Individual injected G0 males were mated to 2-3 *v*^*w*^ females and individual injected G0 females were mated to 2-3 *v*^*w*^ males. Progeny were then subsequently screened for the loss of pigment in the retina. The progeny from individual crosses that resulted in *vermilion*, non-pigmented progeny, were saved. From each positive line, at least two individual *vermilion* progeny were backcrossed to the parental injected strain, either HB or GA-1 to test for genetic drive and any remaining *vermilion* progeny were subjected to PCR and sequence confirmation for CRISPR/Cas9 editing and presence of Cas9.

PCR and Sequence Confirmation of CRISPR/Cas9 editing: Genomic DNA was isolated from individual *Tribolium* by crushing an individual in 50μl of extraction buffer (100mM Tris-HCl, 50mM EDTA, 1% SDS) with a pestle in an Eppendorf tube. The mixture was subjected to a five-minute incubation at 95°C, then chilled on ice. The mixture was then digested with Proteinase K (50 μg/mL) for 1 hour at 55°C, followed by heat inactivation at 95°C for 5 minutes. 200μl of 0.1X TE buffer was added to dilute the sample. Finally, 100μl of the gDNA solution was purified using the Zymo Genomic DNA Clean and Concentrator –10 kit (Zymo Research #ZD4010) following the manufacturer’s instructions. Amplicons spanning the gRNA target sites were amplified from 1 μL of purified gDNA using HotStar PCR Master Mix (Qiagen). Half of each reaction was run on a 1.5% agarose gel, and the other half of the reaction was purified using the Qiaquick Gel Extraction Kit (Qiagen). The purified fragments were submitted to Eurofins Genomics for Sanger sequencing, and the sequences were analyzed using Sequencher (Gene Codes Corp.). The following primers were used to amplify the *vermilion* DNA flanking the targeted gRNA site: 5’-ACCTAAGGTCACGCGGAAGTATCGCATCGT-3’ and 5’-CAGGAGCCTGAACTGCAGGCTCTGGAACCC -3’ and amplify an 806 bp fragment. The vermillion PCR products were sequenced with the following primer: 5’-TATCGCTTTAGTTAGTCTAAA-3’. The following primers were used to detect the presence of Cas9 5’-CTCTAATCGAAACTAATGGGGAAACTGGAG-3’ and 5’-GTTCGTTATCTTCTGGACTACCCTTCAACT-3’ and amplify an 579 bp fragment.

## Results

Utilizing a process based on the mutagenic chain reaction [4], we developed a set of components to target the *vermilion* locus of *Tribolium*. We chose *vermilion* because it has been successfully targeted by CRISPR/Cas9 [21, 27] and loss of function mutations result in an easily scorable visible phenotype, loss of pigment in the adult eye. We generated two homology directed repair (HDR) back bone constructs in which various versions of Cas9 can be inserted. The backbone constructs differ in the number of gRNAs expressed, one versus two, and subsequently different sequences for the right homology arms based upon the position of the second gRNA utilized (Fig 1). The HDR backbones were designed to disrupt the open reading frame of *vermilion*. The single gRNA backbone contained the TcV95 guide RNA [21] and TcV95 has previously been demonstrated to guide Cas9 for genome editing. The two guide RNA backbone contained TcV95 a second gRNA, TcV412; TcV412 had not been tested previously. For Cas9 expression we utilized two different Cas9 expression vectors. The first was the established hsp68Cas9 cassette [24], whereas Cas9 expression is under the control of the core hsp68 gene promoter, presumably active in both the germline and soma of *Tribolium* and does not need a heat pulse for expression [32]. In addition, the Cas9 has the associated 3’ UTR of *hsp68*. The second Cas9 cassette has Cas9 expression controlled by the putative promoter for *Tribolium nanos*. In *Drosophila, nanos* is a germline specific transcript and thus this cassette would potentially limit Cas9 expression to the germline. In addition, the construct has the associated 3’ UTR of *nanos*, which in *Drosophila* is necessary for both localization and translation at the embryonic posterior pole [33, 34]. Overall, four different constructs were injected. For injections, we chose to inject wild-type pigmented *Tribolium* strains, Henderson Black, and GA-1. For the majority of injections, GA-1 was utilized due to an apparent but not quantified greater fecundity; embryos were readily available for injections from the GA-1 stock but not from Henderson Black.

**Fig 1.**
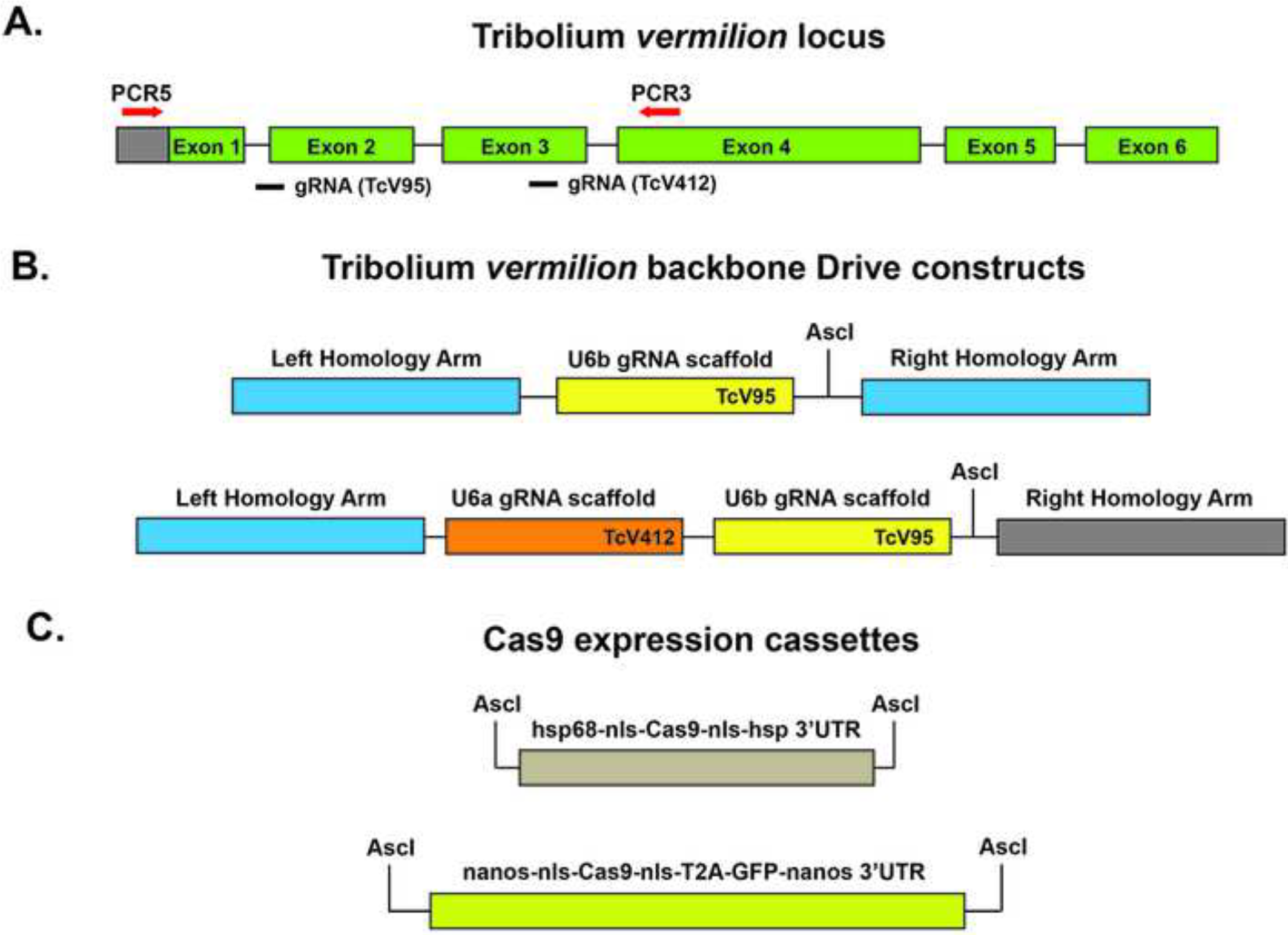
*vermilion* locus and Cas9 homologous recombination drive constructs. A. Schematic of the *vermilion* locus and location of gRNAs and PCR primers for amplifying the region surrounding the potential region of CRISPR/Cas9 editing. B. The backbone constructs for homology directed recombination containing a single gRNA (TcV95) or two different gRNAs (Tcv95 and TcV412). C. Cas9 expression cassettes. Cas9 was either expressed from the ubiquitous *heat shock protein 68 (hsp68)* promoter or potentially the germline restricted *nanos* promoter.

For each construct, at least two rounds of injections were completed. Surviving injected individual males and females were outcrossed to *v*^*w*^ beetles. v^w^ contains a large deletion that removes most of the *vermilion* locus and results in non-pigmented eyes [21]. As such, in the F1 generation potential HDR and drive candidates were identified by the loss of pigment in the retina. For all four constructs, non-pigmented F1 progeny were identified (Table 1). In any particular cross that generated non-pigmented progeny, the number of F1 *vermilion* progeny ranged from a few to over 50%.

**Table 1:**
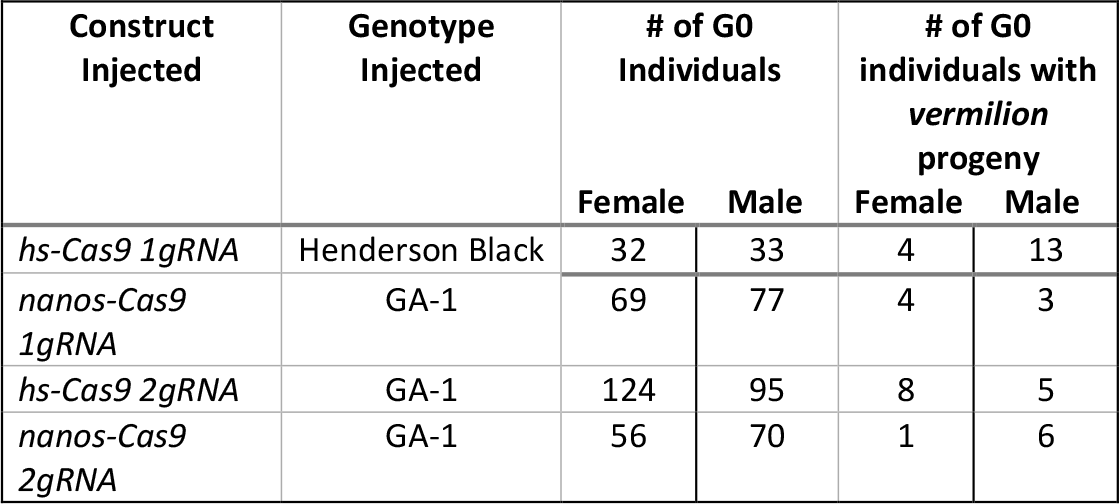
Summary of injections and recovered *vermilion* edited progeny.

Upon the identification of *vermilion* progeny, two subsequent analyses were conducted. To test for the presence of homologous recombination and subsequent functional drive, at least two and preferably a male and virgin *vermilion* F1 progeny were backcrossed to GA-1 pigmented individuals. If homologous recombination repair did occur and genetic drive was active, we would expect 50% of the resulting progeny to contain non-pigmented retinas, the result of inheriting the CRISPR edited allele, and 50% pigmented, the result of inheriting the v^w^ allele and the presence of a wild type *vermilion* allele from the GA-1 stock. None of the presumed CRISPR edited *vermilion* progeny when backcrossed to GA-1 resulted in the presence of *vermilion* progeny. To eliminate the possibility that HDR did occur but the construct failed to promote genetic drive, genomic DNA was isolated from original F1 *vermilion* progeny, derived from the injected G0 individuals, and assayed for the presence Cas9. We did not detect the presence of a Cas9 PCR amplicon in any of the samples. These two results combined suggested that CRISPR/Cas9 directed HDR did not occur.

Lastly, we needed to confirm that the *vermilion* progeny obtained were the result of CRISPR/Cas9 editing and with respect to the two gRNA constructs, do both gRNAs work and can we detect editing with both gRNAs on the same DNA molecule? For each F1 progeny from the G0 injected individuals that resulted in *vermilion*, non-pigmented retinas, genomic DNA was isolated and primers were used to amplify DNA spanning across the location of the directed gRNA cuts (Fig 1). For those samples that resulted in a PCR product (Fig 2A and 3A), the PCR product was sequenced and examined for changes in the DNA. As expected, the one gRNA backbone HDR construct with either Cas9 under the control of *hsp68* or *nanos* resulted in deletions, insertions or a combination of both (Fig 2B). With respect to the 2 gRNA two gRNA backbone HDR construct, we found examples in which a single gRNA was utilized and examples of DNA molecules that contained evidence of CRISPR/Cas9 editing at both targeted sites on the same DNA molecule with either *hsp68* or *nanos* promoter driving Cas9 expression (Fig 3B). Furthermore, we recovered one edited mutation, f12, that deleted the entire region, except for the insertion of 6 nucleotides, between the two gRNA target sites (Fig 3B). Overall, our results demonstrated that both HDR backbone constructs, gRNAs, and Cas9 expression cassettes are capable of inducing double strand breaks and the lack of homology directed repair was not necessarily due to the functionality of the designed components.

**Fig 2.**
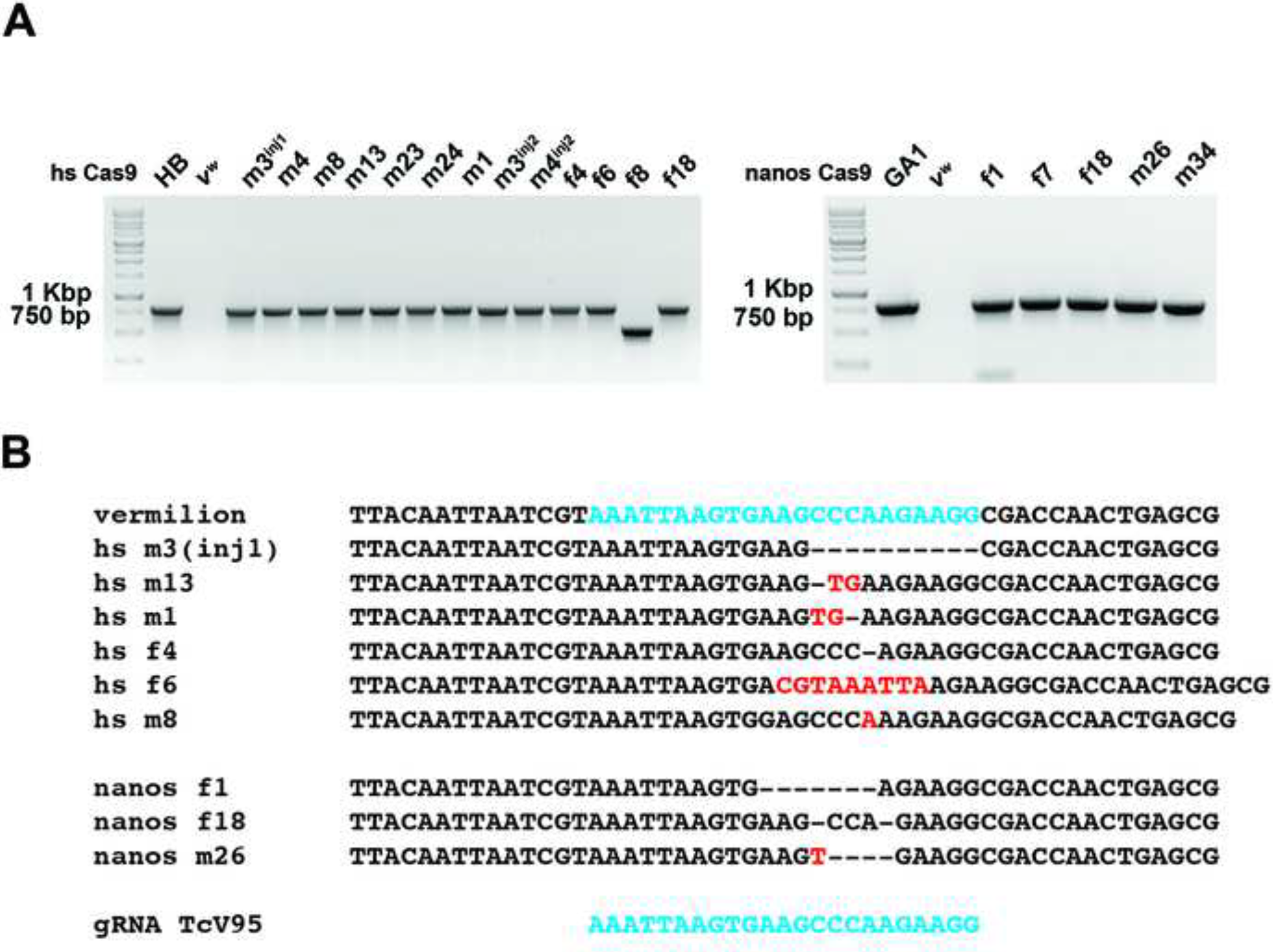
PCR and Sequencing results of CRISPR/Cas9 editing utilizing a single gRNA combined with Cas9 expressed from the *hsp68* or *nanos* promoters. A. PCR amplicons from isolated vermilion progeny from injections HDR backbones containing one gRNA and Cas9. The PCR amplicons are relatively equal in size or smaller to the expected to the unedited *vermilion* locus, demonstrating the lack of homologous recombination in the *vermilion* progeny. The numbers above each lane refer to the GO injected individual, m-male and f-female. B. Alignment and sequence confirmation and identification and nature of CRISPR/Cas9 editing in a sample of the PCR amplicons from the isolated *vermilion* progeny. Dashes represent deleted bases and red color bases indicate base pair insertions.

**Fig 3.**
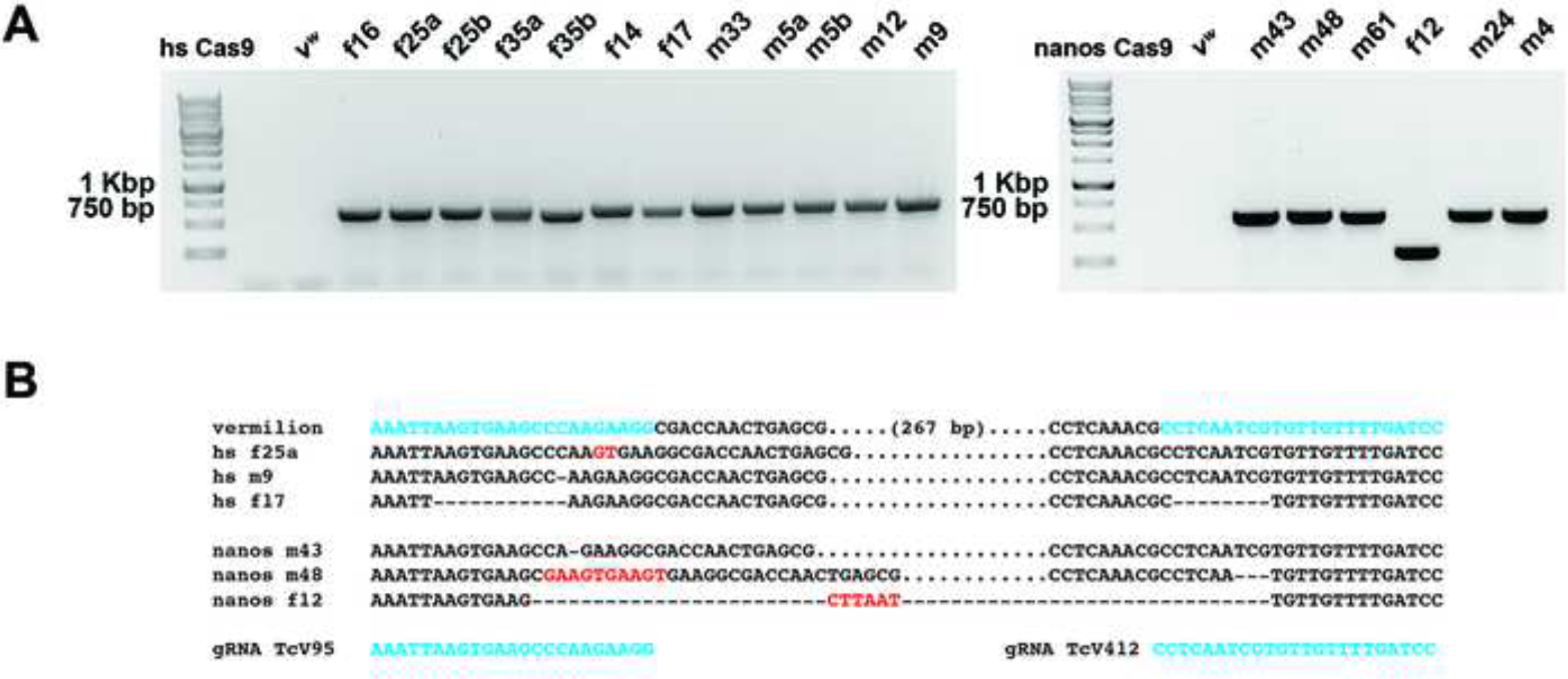
PCR and Sequencing results of CRISPR/Cas9 editing utilizing a two gRNAs combined with Cas9 expressed from the *hsp68* or *nanos* promoters. A. PCR amplicons from isolated *vermilion* progeny from injections HDR backbones containing two gRNAs and Cas9. The PCR amplicons are relatively equal in size or smaller to the expected to the unedited *vermilion* locus, demonstrating the lack of homologous recombination in the *vermilion* progeny. The numbers above each lane refer to the GO injected individual, m-male and f-female. B. Alignment and sequence confirmation and identification and nature of CRISPR/Cas9 editing in a sample of the PCR amplicons from the isolated *vermilion* progeny. Dashes represent deleted bases. Red color bases indicate base pair insertions and a dot represents base pairs that are present but not shown.

## Discussion

The possibilities for failure of HDR and subsequent genetic drive are many, but we have demonstrated a few key principles that will aid future editing in *Tribolium*. We have demonstrated the functionality of our gRNAs and the ability to direct Cas9 editing to two sites on the same DNA molecule. In addition, we have shown the functionality of our *nanos* Cas9 cassette, providing a second methodology to express Cas9. Furthermore, we also noted a high efficiency of recovering CRISPR/Cas9 edited alleles of *vermilion* in our experiments, 8% of surviving G0 progeny. Even though an unbiased study would be required to confirm this observation, we feel the inclusion of both the gRNAs and the expression cassette for Cas9 on the same plasmid was responsible for an increase in efficiency. Our end goal was to achieve homologous directed repair and subsequent test for genetic drive, however our results demonstrated non-homology end joining (NHEJ) was the preferred method for repair.

Is there a possible methodology to skew repair towards homology directed repair versus NHEJ in future work? One possibility would be to eliminate the function of Lig4; Lig4 is a critical enzyme for NHEJ [35, 36]. In *Drosophila, Lig4* mutants are homozygous viable [37] and editing of the genome with zinc-finger nucleases or CRISPR/Cas9 viable homology directed repair was biased towards HDR [38, 39]. In addition, RNAi depletion of Lig4 in *Drosophila* tissue culture cells increased the frequency of HDR versus NHEJ when utilizing CRISPR/Cas9 [40].

Interestingly, *Tribolium* has not one but two orthologs of *Lig4* (LOC657210 and LOC657043) and thus future experiments will require mutating of one or both and test whether NHEJ with respect to CRISPR/Cas9 editing is decreased with subsequent increase in the frequency of HDR. Moreover, in *Drosophila* HDR frequencies were the greatest when plasmids encoding the gRNAs and the homology repair template were injected together into *Drosophila* transgenic lines expressing Cas9 in the germline [39]. Our results here now demonstrate the possibility of using the *nanos* promoter to achieve germline expression. And, although Cas9 expression may not be limited to the germline, a transgenic version of the hs-Cas9 cassette exists [27] and can be used in future CRISPR/Cas9 HDR attempts.

## Acknowledgments

This work was supported by NIFA/USDA 2019-33522-30064 (M.J.W, A.C.Z., G.E.Z.) and NSF grant IOS-1928781 (A.C.Z.).

## Data Availability

All relevant data are within the paper and its Supporting Information files. All vectors generated are available through the Drosophila Genomics Resource Center at Indiana University and complete sequences of vectors can be found in Supplement Files.

